# Nitric Oxide Mediates Neuro-Glial Interaction that Shapes *Drosophila* Circadian Behavior

**DOI:** 10.1101/700971

**Authors:** Anatoly Kozlov, Emi Nagoshi

**Affiliations:** Department of Genetics and Evolution, Sciences III, University of Geneva, 30 Quai Ernest-Ansermet, Geneva-4, CH-1211, Switzerland

## Abstract

*Drosophila* circadian behavior relies on the network of heterogeneous groups of clock neurons. Short -and long-range signaling within the pacemaker circuit coordinates molecular and neural rhythms of clock neurons to generate coherent behavioral output. The neurochemistry of circadian behavior is complex and remains incompletely understood. Here we demonstrate that the gaseous messenger nitric oxide (NO) is a signaling molecule linking circadian pacemaker to rhythmic locomotor activity. We show that two independent mutants lacking nitric oxide synthase (NOS) have severely disturbed locomotor behavior both in light-dark cycles and constant darkness, although molecular clocks in the main pacemaker neurons are unaffected. Behavioral phenotypes are due in part to the malformation of neurites of the main pacemaker neurons, s-LNvs. Using cell-type selective and stage-specific gain -and loss-of-function of NOS, we demonstrate that NO secreted from diverse cellular clusters non-cell-autonomously affect molecular and behavioral rhythms. We further identify glia as a major source of NO that regulates circadian locomotor output. These results reveal for the first time the critical role of NO signaling in *the Drosophila* circadian system and highlight the importance of neuro-glial interaction in the neural circuit output.

**Author summary:** Circadian rhythms are daily cycles of physiological and behavioral processes found in most plants and animals on our planet from cyanobacteria to humans. Circadian rhythms allow organisms to anticipate routine daily and annual changes of environmental conditions and efficiently adapt to them. Fruit fly *Drosophila melanogaster* is an excellent model to study this phenomenon, as its versatile toolkit enables the study of genetic, molecular and neuronal mechanisms of rhythm generation. Here we report for the first time that gasotransmitter nitric oxide (NO) has a broad, multi-faceted impact on *Drosophila* circadian rhythms, which takes place both during the development and the adulthood. We also show that one of the important contributors of NO to circadian rhythms are glial cells. The second finding highlights that circadian rhythms of higher organisms are not simply controlled by the small number of pacemaker neurons but are generated by the system that consists of many different players, including glia.

## Introduction

Our environment undergoes daily fluctuations in solar illumination, temperature, and other parameters. Organisms across the phylogenetic tree are equipped with circadian clocks, which help predict daily environmental changes and create temporal patterns of behavioral and physiological processes in concordance with the environmental cycle. *Drosophila melanogaster* remains a handy model to study this phenomenon ever since Konopka and Benzer identified the first clock gene, *period*, in this organism (1).

*Drosophila* circadian clocks rely on transcriptional-translational feedback loops that operate using an evolutionarily conserved principle. In the main loop, CLOCK/CYCLE (CLK/CYC) heterodimers bind to the E-boxes in the promoter regions of the *period* (*per*) and *timeless* (*tim*) genes and activate their transcription. PER and TIM proteins undergo post-translational modifications and enter the nucleus to suppress their own production by inhibiting CLK/CYC activity. CLK/CYC also activates transcription of the genes encoding the basic-zipper regulators PAR DOMAIN PROTEIN 1 (PDP-1) and VRILLE (VRI), which activates and inhibits *Clk* gene expression, respectively. Thus, positive -and negative-feedback loops created by PDP-1 and VRI with CLK/CYC are interlocked with the main negative-feedback loop and ensure the generation of 24-h rhythms (2,3).

In the fly brain, molecular clocks are present in ca.150 so-called clock neurons, which form the pacemaker circuit controlling circadian behavior. Clock neurons are classified into groups according to their morphological characteristics and location: small and large lateral ventral neurons (s -and l-LNvs), lateral dorsal neurons (LNds), lateral posterior neurons (LPNs) and three groups of dorsal neurons (DN1s, DN2s, DN3s) (4,5). Although all clock neurons express a common set of clock genes, they are heterogeneous in terms of neurotransmitter/neuropeptide phenotype, function, and composition of the molecular clock. Neuropeptide pigment-dispersing factor (PDF) is uniquely secreted from the l-LNvs and 4 out of 5 s-LNvs. Several other neuropeptides, including small neuropeptide F (sNPF) and ion transport peptide (ITP), and classical neurotransmitters such as glutamate and glycine, are also expressed across pacemaker circuit (6,7). PDF-positive s-LNvs are designated as the Morning (M) oscillator, whereas LNds together with the PDF-negative 5th s-LNv consist of the Evening (E) oscillator. Under the light-dark (LD) experimental conditions, the M and E oscillators drive the morning and evening anticipatory increments of locomotor activity, respectively. The M oscillator is also the master pacemaker of the free-running locomotor rhythms in constant darkness (DD) (8–10).

Neuropeptide PDF as well as the unique composition and regulatory mechanisms of the molecular clock underlie the distinct role of the M oscillator. The main negative-feedback loop of the M oscillator’s molecular clock employs a specific phosphorylation program that regulates the nuclear translocation of PER/TIM complex (11). The nuclear receptor UNFULFILLED (UNF) is almost uniquely present in the lateral neurons within the circadian circuit (12,13). UNF accumulates rhythmically in the s-LNvs and, in cooperation with another nuclear receptor E75, enhances CLK-dependent *per* transcription. Thus, UNF and E75 consist a positive limb of an additional feedback loop in specific to the s-LNv molecular clock. Because UNF and E75 also play critical roles in the development of the s-LNvs, knockdown of either gene during development or adulthood results in low rhythmicity and extended period, respectively (12,14).

Nuclear receptors (NRs) are a superfamily of proteins that function as ligand-dependent transcriptional regulators (15). The ligands are small lipophilic molecules that can diffuse across the cell membrane, such as thyroid and steroid hormones. In *Drosophila melanogaster*, only two lipophilic hormones, 20-hydroxyecdyson (20E) and the sesquiterpenoid juvenile hormone (JH) are known nuclear receptor ligands, which have critical roles in developmental processes, including molting, puparium formation, and neurogenesis (15–17). Although many NRs remain orphan without a known ligand, diatomic gases nitric oxide (NO) and carbon monoxide (CO) can bind and regulate the activity of some NRs. Several studies have demonstrated *in vitro* and *in vivo* that NO binds to E75 and regulates its interaction with DHR3 (18), SMRTER (19), and UNF (20) in different tissues during development. Thus, the binding of NO to E75 confers an important switching mechanism in various developmental processes.

NO is an unconventional messenger involved in numerous biological functions, including immune defense, respiration, intracellular signaling and neurotransmission (21,22). NO can act locally near the source of its production. It can diffuse across membranes and also act as a long-range signaling molecule (22,23). NO signaling is broadly classified into the classical pathway mediated by cGMP and cGMP-independent non-classical one involving diverse mechanisms such as posttranslational modifications and transcriptional regulations (22,24). In mammals, the importance of NO signaling in the light-dependent phase-resetting and maintenance of rhythmicity (25–28) is established. These effects were largely explained by the canonical NO/cGMP signaling (29–31). However, whether NO has a regulatory role in *Drosophila* circadian behavior has never been addressed.

Here we explore the role of NO in circadian locomotor behavior of *Drosophila* using multiple genetic approaches. We present evidence that NO signaling is necessary for both light-dependent and free-running circadian behavior. NO acts cell-autonomously as well as non-cell-autonomously at multiple processes required for generating behavior, including axonal morphogenesis, pacing of molecular clocks and output control. We identify glial cells as a major source of NO that controls free-running locomotor output. Our results highlight the complexity of locomotor behavior regulation and oft-neglected importance of glia in the regulation of behavior.

## Results

### *dNOS* deletion mutants show abnormal circadian behavior

NO is chiefly produced by an enzyme nitric oxide synthase (NOS) through the conversion of arginine into citrulline using NADPH as a cofactor (32,33). Three distinct NOS isoforms (endothelial e-NOS, inducible i-NOS, and neuronal n-NOS) exist in mammals, whereas *Drosophila* has a single *NOS* (*dNOS*) gene that produces 10 splice variants (Fig. S1) (34). Since NOS functions as homodimers, alternatively spliced variants, most of which encode truncated proteins, are proposed to act as dominant negatives (35). To investigate the possible roles of NO in fly circadian rhythms, we took advantage of two *NOS* CRISPR deletion mutants, *NOS Δ all*, and *NOS Δ ter* (20). The former has a deletion of the entire NOS locus, while the latter is a partial deletion mutant lacking exons 1 to 6 but bears intact two uncharacterized genes within the *NOS* locus (Fig 1A). RT-qPCR using the primers targeting the exons commonly included in all variants (exon 10 and 11) confirmed the absence of the full-length NOS1 mRNA expression in *NOS Δ all* mutants (Fig. 1B) A reduced level of the product was detected in *NOS Δ ter* mutants, consistent with the location of the deletion. Furthermore, we directly measured NO production in cultured whole brains using a fluorescent dye DAR4-M (36). NO was virtually undetectable in both *NOS Δ all* and *NOS Δ ter* strains in this assay, confirming that both are complete loss-of-function mutants (Fig. 1C).

**Figure 1.**
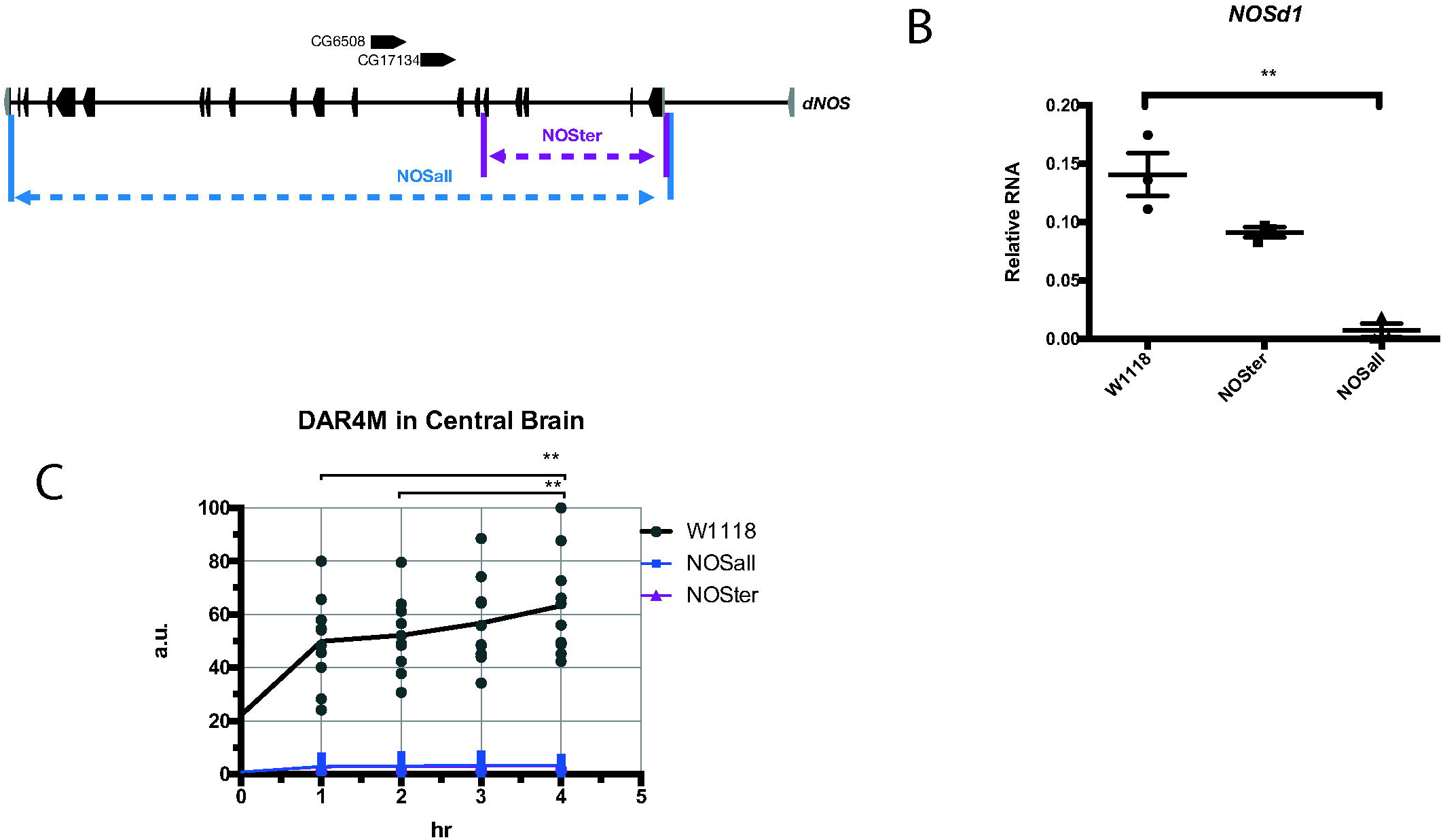
*NOSΔ* mutants do not produce nitric oxide. (A) *NOS* gene, *NOSΔ*all, and *NOSΔ ter* mutants and two reverse genes residing within the *NOS* locus. (B) mRNA levels of *NOS1*, a full-length functional isoform of NOS (NOSd1), at ZT2 in the heads of *NOS Δ* mutants and *w*^*1118*^ were analyzed using qPCR.*\*\*p*<0.01 (Student’s test). (C) NO levels were measured using DAR4-M dye in the brains of *NOS Δ* mutants and *w*^*1118*^ in brain explants for 4 h using timelapse microscopy. *\*\*p*<0.01 (Student’s test)

Having validated the deletion mutants, we next tested their locomotor activities in LD and DD paradigms. Homozygous mutants had strongly reduced rhythmicity in DD. Trans-heterozygous of two deletion alleles was equally detrimental to DD rhythmicity, whereas heterozygous mutations had no effect on rhythmicity (Table 1 and Fig. 2A). Moreover, morning activity patterns in LD were strongly impaired in homozygous and trans-heterozygous mutants, nearly lacking both anticipatory increments of activity before lights-on and the startling reaction to light (Fig. 2B). The severe reduction of the startling response to light suggests an impairment in photoreception through the compound eyes known to be necessary for this phenomenon (37,38). This is consistent with the fact that NO/cGMP signaling is necessary for neurites patterning of the receptor neurons in the optic lobe during development (39). On the other hand, low rhythmicity in DD and poor morning anticipation are indicative of the dysfunction of the s-LNvs, the Master and Morning oscillator.

**Table 1.** Free-running locomotor rhythms.

| Temperature | Genotype | Period ± SEM (hr) | Power ± SEM | n | %R |
| --- | --- | --- | --- | --- | --- |
| 25°C | *W1118* | 23.5±0.05 | 167.5±9.3 | 124 | 93.6 |
| | *CantonS* | 24.1±0.35 | 102.3±18.0 | 27 | 84.4 |
| | *NOSter/+* | 23.7±0.04 | 217.7±10.5 | 127 | 91.3 |
| | *NOSter* | 23.2±0.2 | 84.3±29.0 | 105 | 21.0*** |
| | *NOSall/+* | 23.7±0.04 | 245.1±12.1 | 126 | 76.9 |
| | *NOSall* | 23.6±0.06 | 149.4±15.5 | 173 | 56.1** |
| | *NOSall/NOSter* | 23.5±0.2 | 79.0±17.4 | 59 | 66.1 |
| macNOS oxp | *>macNOS* | 23.4±0.05 | 59.4±2.1 | 89 | 85.4 |
| LNvs | *PDF>macNOS* | 24.5±0.15*** | 51.7±6.1 | 25 | 76.6 |
| s-LNVs | *R6>macNOS* | 23.8±0.09 | 69.9±5.2 | 31 | 65.5** |
| MB | *D52H>macNOS* | 24.2±0.05 | 145.2±13.9 | 28 | 92.9 |
| Clock neurons | *tim>macNOS* | 24.7±0.1**** | 61.3±6.3 | 30 | 36.7**** |
| Clock neurons | *Clk1982>macNOS* | 23.9±0.05 | 85.5±7.6 | 30 | 76.7 |
| OL | *GMR33H10>macNOS* | 23.7±0.09 | 85.2±8.7 | 62 | 53.2**** |
| OL | *GMR79D04>macNOS* | 23.9±0.09 | 43.3±4.9 | 63 | 28.6**** |
| OL | *GMR85B12>macNOS* | 23.6±0.1 | 53.1±7.9 | 60 | 61.7*** |
| Glia | *Repo>macNOS* | 23.5±0.05 | 98.2±5.0 | 28 | 29.8**** |
| Pan-neuronal | *GMR57C10>macNOS* | 25.0±0.4* | 65.9±12.9 | 61 | 32.8**** |
| Pan-neuronal | *Elav>macNOS* | 23.7±0.05 | 83.5±6.4 | 59 | 81.4 |
| NOS KD | *NOS-RNAi<sup>27725</sup>/+* | 23.7±0.04 | 110.7±0.04 | 145 | 84.0 |
| LNvs | *PDF>NOS-RNAi* | 24.4±0.03 | 184.1±13.9 | 26 | 100 |
| s-LNVs | *R6>NOS-RNAi* | 23.6±0.04 | 113.0±5.5 | 60 | 80 |
| MB | *D52H>NOS-RNAi* | 23.6±0.4 | 105.4±10.5 | 29 | 65.5* |
| MB | *OK107>NOS-RNAi* | 23.5±0.05 | 141.2±12.0 | 32 | 96.9 |
| Photoreceptors | *GMR>NOS-RNAi* | 23.6±0.06 | 200.1±12.3 | 60 | 95.2 |
| Clock neurons | *tim>NOS-RNAi* | 24.4±0.2 | 166.2±10.4 | 47 | 38.3**** |
| Clock neurons | *Clk1982>NOS-RNAi* | 23.6±0.02 | 173.3±5.03 | 62 | 88.7 |
| OL | *GMR33H10>NOS-RNAi* | 23.5±0.03 | 164±7.1 | 32 | 96.9 |
| OL | *GMR79D04 >NOS-RNAi* | 25.1±0.2** | 105.2±7.9 | 59 | 61.0** |
| OL | *GMR85B12 >NOS-RNAi* | 23.6±0.03 | 121.3±6.9 | 44 | 70.5** |
| Glia | *Repo>NOS-RNAi* | 23.6±0.2 | 79.4±6.6 | 63 | 32.1**** |
| Pan-neuronal | *GMR57C10>NOS-RNAi* | 26.2±0.2**** | 121.2±8.2 | 60 | 55.0** |
| Pan-neuronal | *elav>NOS-RNAi* | 23.5±0.05 | 122.3±9.1 | 95 | 76.8 |
| CTR | *PDF>* | 24.1±0.04 | 132.7±6.9 | 58 | 96.8 |
| | *R6>* | 23.4±0.05 | 105.2±5.7 | 45 | 85.1 |
| | *D52H>* | 23.9±0.04 | 177.3±8.3 | 10 | 90.5 |
| | *OK107>* | 23.7±0.06 | 139.6±8.5 | 30 | 96.8 |
| | *GMR* | 23.6±0.03 | 187.5±6.1 | 28 | 93.3 |
| | *Tim>* | 24.1±0.05 | 125.4±6.3 | 82 | 90.2 |
| | *Clk1982>* | 23.6±0.03 | 150.6±7.4 | 60 | 86.7 |
|  | *GMR33H10>* | 23.1±0.05 | 144.9±6.2 | 60 | 85.0 |
|  | *GMR79D04 >* | 24.3±0.2 | 96.6±5.5 | 64 | 84.4 |
|  | *GMR85B12 >* | 23.5±0.04 | 128.5±6.9 | 91 | 89.0 |
|  | *Repo>* | 23.3±0.05 | 109.2±6.9 | 83 | 82.0 |
|  | *GMR57C10>* | 24.2±0.2 | 117.9±4.9 | 60 | 78.3 |
|  | *Elav>* | 23.7±0.05 | 143.0±7.8 | 120 | 82.8 |
| **18°C ->29°C** | *Repo>NOS-RNAi* | 24.1±0.5 | 47.2±7.3 | 21 | 47.6* |
| **Adult-only KD** | *GMR79D04>NOS-RNAi* | 24.7±0.3 | 93.1±8.3 | 28 | 75.0 |
|  | *GMR85B12>NOS-RNAi* | 25.6±0.3 | 86.0±12.5 | 17 | 56.7 |
|  | *GMR57C10>NOS-RNAi* | 26.8±0.07*** | 165.3±11.5 | 25 | 96.0 |
| **CTR** | *NOS-RNAi/Tub-Gal80ts* | 25.6±0.3 | 89.2±6.5 | 29 | 79.3 |
|  | *Repo>* | 23.5±0.6 | 114.2±9.3 | 29 | 72.4 |
|  | *GMR79D04>* | 23.3±0.07 | 76.0±8.7 | 28 | 67.9 |
|  | *GMR85B12>* | 23.2±0.09 | 74.2±15.1 | 28 | 32.1 |
|  | *GMR57C10>* | 23.2±0.06 | 121.7±9.74 | 30 | 86.7 |

$a_n$ , Number of flies; %R, % of rhythmic flies. Rhythmicity was compared with the driver control or with control genotypes (*CantonS* for *NOS* mutants) by a $\chi^2$ test. Periods were compared with the controls (Student's *t* test): \**p* 0.05; \*\**p* 0.01; \*\*\**p* 0.001; \*\*\*\**p* 0.0001.

**Figure 2.**
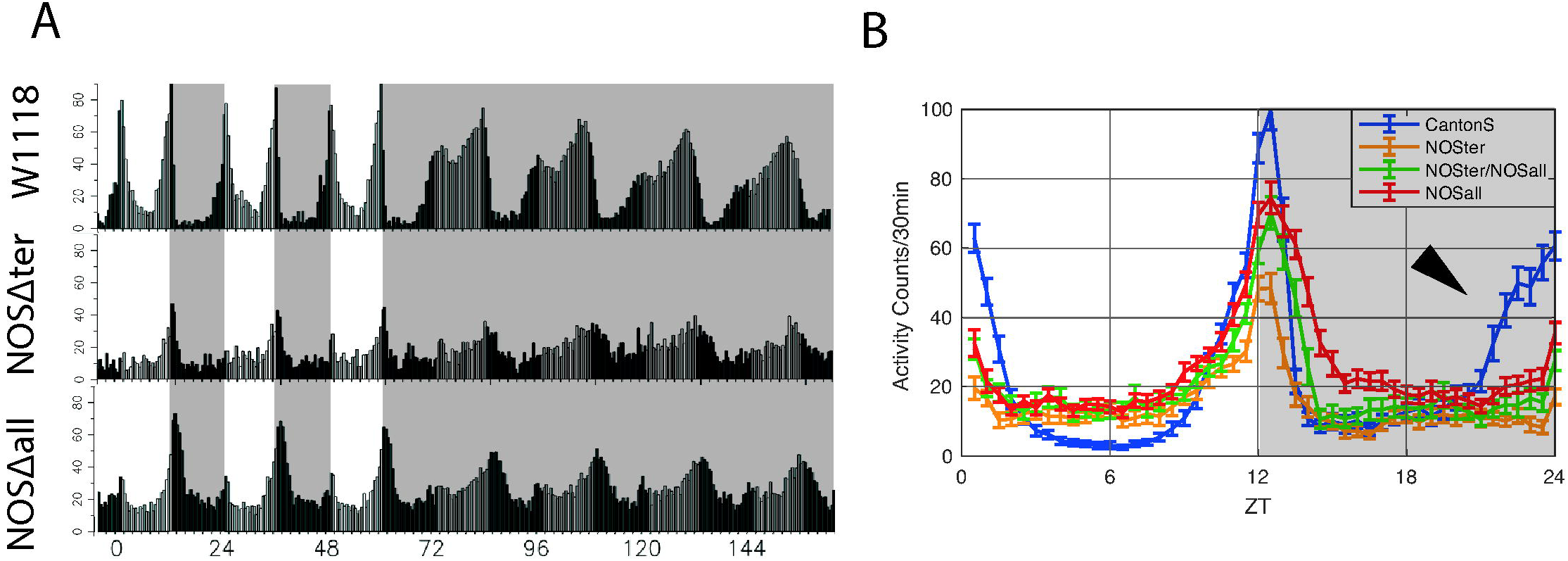
*NOS* deletion impairs the circadian behavior in light-dark cycles and constant darkness. (A) Locomotor activity histograms for group average activity of *w*^*1118*^ and *NOS Δ* mutants. (N=32)(B) Histograms of group average activities of *w*^*1118*^ and *NOS Δ* mutants in LD, an average of 3 days.

### NOS is a regulator of morphogenesis of the s-LNv axons

NO signaling plays critical roles in various developmental processes in the nervous system, including neurite patterning of the visual system and axon pruning/regrowth of mushroom body (MB) neurons (18–20,39,40). Therefore, to explore the possible effect of *NOS* loss-of-function on the development of the s-LNvs, we expressed a membrane-targeted yellow fluorescent protein mCD8::VENUS with the *gal1118-Gal4* driver and visually inspected s-LNv axonal morphology. To our surprise, normally a very orderly branching pattern of terminal neurites was severely disturbed in *NOS* homozygous mutants (Fig. 3A). The neurites were extended in length and branching pattern was highly disordered and fuzzy (Fig. 3A and B). Circadian change of axonal termini (41) was not evident in mutants due to the highly disordered structure. This disorderly morphology is reminiscent of the axon pruning defects of MB neurons in *NOS* deletion mutants (20). Since NO promotes MB neuron axon pruning and degeneration by inhibiting UNF/E75 heterodimer formation (20), we wondered if similar mechanisms are involved in the structural maturation of the s-LNvs. To probe this idea, we visualized s-LNv projections by expressing mCD8::VENUS while constitutively knocking down UNF. We found that s-LNv axonal morphology was affected in a similar manner in this condition, having more numerous and disordered neurites than the control (Suppl. Fig S2). These results suggest that NOS is necessary for axonal morphogenesis of the s-LNvs and part of its role may be via controlling the activity of UNF.

**Figure 3.**
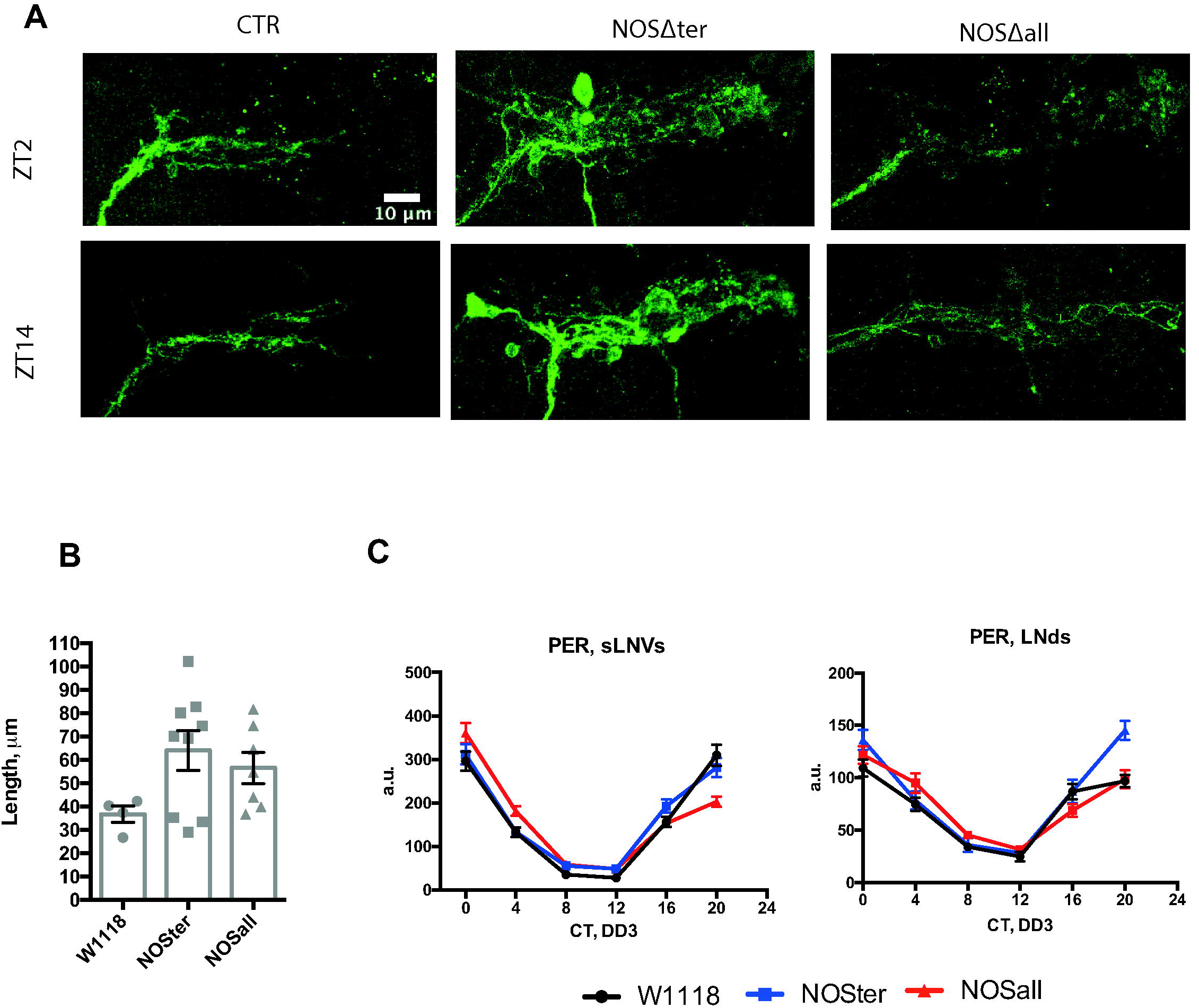
*NOS* deletion causes the malformation of the axons of the s-LNvs but does not affect their molecular clocks. (A) s-LNv axonal terminal projections were visualized by expressing *mCD8::Venus* and staining with anti-GFP antibodies at ZT2 and ZT14. Representative confocal images are shown. (B) Quantifications of the length of the terminal branches at ZT2. (C) PER levels of *w*^*1118*^ and *NOS Δ* mutants in the s-LNvs and LNds analyzed every 4 h on DD3. NOS deletion does not affect PER rhythms.

To better understand the nature of the low rhythmicity in *NOS* mutants, we also performed around-the-clock immunostaining of a key clock component PER on the third day of constant darkness (DD3). Neither the phase nor the amplitude of the molecular clocks of the s-LNvs was affected in *NOS* mutants. Molecular clocks of the LNds also maintained high-amplitude 24-h rhythms in mutants (Fig. 3C). Therefore, the arrhythmic behavioral phenotype of *NOS Δ* mutants is uncoupled from the state of the molecular clocks and caused by the developmental impairments. It probably entails the wrong synaptic connectivity of the malformed s-LNvs axonal terminals, in addition to defects in possibly many other cells involved in locomotor output. Malformation of s-LNv axons is likely to be also responsible for lack of morning anticipatory behavior.

### NO from diverse cellular groups can modulate the state of the molecular clocks and behavioral output

Whereas NOS is undoubtedly important for developmental processes, the fact that NO continues to be produced in the brains of adult flies (Fig. 1C) suggests that NO may also have a post-developmental role in regulating circadian rhythms. It was previously shown using an anti-NOS serum (42) that NOS is expressed almost everywhere in the brain. However, the anti-NOS serum does not distinguish various NOS isoforms and thus the picture may not be identical to the loci of active NO production. Separately, classical histochemical studies of NADPH diaphorase activity of NOS and soluble guanylate cyclase (sGC)/cGMP immunohistochemistry have suggested that NOS is active in sensory pathways including visual system, in memory circuits including the calyx of mushroom body, in the central complex and also in some glial cells (32,39,43–45). Since these were all indirect assessments of NO production, we analyzed the localization of NO in the brain using the NO-specific fluorescent probe DAR4-M. DAR4-M staining showed distinct patterns of cell bodies and neurites in many areas. The signal was particularly high within and around the central complex and in the optic lobe, with cell bodies arranged in concentric semicircles reminiscent of the laminar and medulla glial cells (46) (Figure 4A). These patterns were overall similar to those described for the localization of NADPH diaphorase activity and cGC/cGMP. In addition, since DAR4-M staining is not sensitive enough to assess daily variation of NO production, we examined the temporal expression pattern of the functional isoform *dNOS1* using qPCR. We found that mRNA of the *dNOS1* was rhythmically expressed in the fly head, peaking around ZT10 in LD (Fig. 4B), suggesting that overall NO levels in the brain may be circadian.

**Figure 4.**
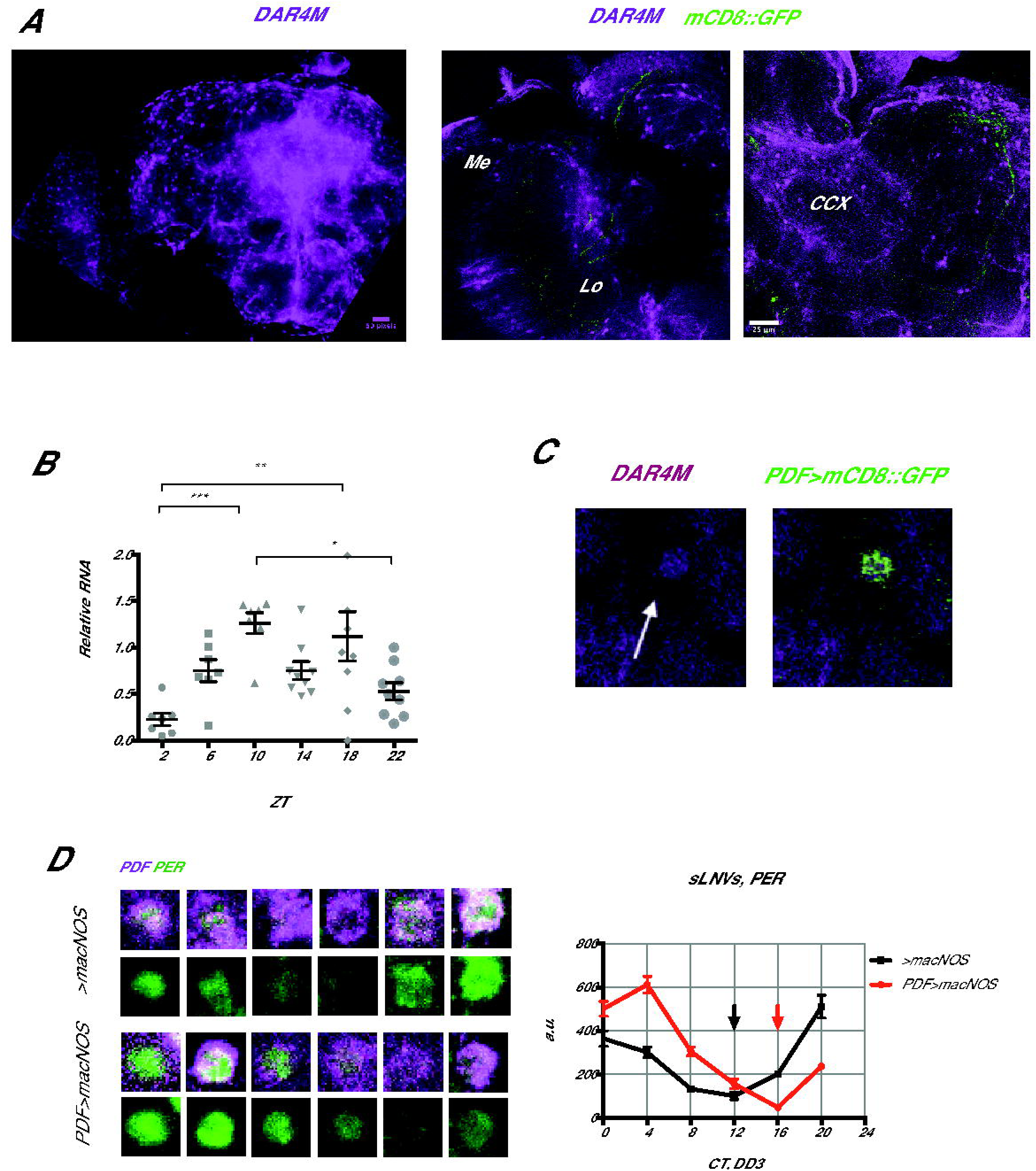
NO production in the brain and its effect on the s-LNv molecular clocks. (A) Intracellular NO staining with DAR4-M staining showed an accumulation of NO in the optic lobe (OL) and in and around the central complex (CCX) of *w*^*1118*^ flies. (B) Around-the-clock measurement of the *dNOS1* isoform mRNA levels by qPCR in LD. *\*p*<0.05, *\*\*p*<0.01, *\*\*\*p*<0.001 (one-way ANOVA test) (C) Enrichment of the DAR4-M signal within the s-LNv cell body. (D) PER levels of *w*^*1118*^ and *PDF>macNOS* mutants were measured every 4 h on DD3. Red and black arrows point at the trough of PER rhythms in *Pdf* > *macNOS* and control flies, respectively. PER rhythms are delayed by approximately 4 hours in *PDF>macNOS* flies.

Interestingly, we detected a slight enrichment of NO-positive fluorescence within the s-LNvs marked with *gal1118 > mCD8::Venus* (Fig. 4C). This was unexpected because previous transcriptome studies found very little or no *NOS* expression within the s-LNvs (13,47,48). This suggests that NO produced elsewhere migrates to the s-LNvs. Within the s-LNvs, transcriptional regulation by E75 and UNF is a unique and important node of the molecular clockwork (12,14). Intriguingly, it was shown that heterodimerization of E75 and UNF is controlled by NO *in vivo* and *in vitro* (20). Therefore, we asked whether the state of the molecular clocks can be modulated by increasing NO within the s-LNvs.

To this end, we overexpressed a macrophage-derived constitutively active NOS (macNOS) under the UAS control (18) using *Pdf-Gal4* and performed PER staining on DD3 every 4 h (Fig. 4D). The increase of NO within the PDF positive neurons was confirmed by DAR4-M staining (Suppl. Fig. S3). This manipulation lead to a delay of the PER induction phase by about 4 h without dampening the amplitude of PER rhythms. The results are consistent with the known role of UNF and E75 and regulation of their heterodimerization with NO: high levels of NO disrupt UNF/E75 dimer and delays PER rising phase that is normally enhanced by UNF/E75.

*Pdf>macNOS* flies had a slight extension of free-running period and the reduction in rhythmicity. But the reduced rhythmicity was not statistically significant compared with the driver control. Because *Pdf-GAL4* is expressed in both s -and l-LNvs, these phenotypes may be a compound effect from both cell types. When we expressed macNOS under the s-LNv-specific *R6-Gal4* driver, we observed a reduction in rhythmicity but no differences in the period length (Table 1). Neither manipulation consistently affected behavioral patterns in LD (Suppl. Fig.S4). Therefore, the major behavioral consequence of forced NO production in the s-LNvs is a reduction in free-running rhythmicity. This is probably a consequence of the misalignment of molecular phases between the s-LNvs and other clock neurons (Fig. 4D).

The results of the DAR4-M staining and the finding that NO can regulate the state of the molecular clocks prompted us to investigate whether NO produced in specific cell types or brain area is important for normal circadian locomotor activity. Therefore, taking into account the notion that NO can act both locally and remotely, we selected a set of GAL4 drivers and drove the expression of macNOS. Locomotor activity of these flies was assayed in standard LD-DD conditions.

As summarized in Table 1, we used two clock cell-specific drivers *Tim-GAL4* and *Clk1982-GAL4*; a mushroom body-specific driver *DH52-GAL4*; three generic optic lobe-specific drivers *GMR33H10-GAL4, GMR79D04GAL4-*, and *GMR85B12-GAL4 (Suppl. Table S1)*; a glia-specific driver *Repo-GAL4*; and two pan-neuronal drivers *elav-GAL4* and *R57C10-GAL4*. Strikingly, all of them except *elav-GAL4* induced a reduction of rhythmicity upon overexpression of macNOS. This is most likely because pan-neuronal *elav-GAL4* is a weaker driver than another pan-neuronal driver *R57C10*-*GAL4*. The reduction of rhythmicity was markedly dramatic with *Repo, tim* and optic lobe specific drivers. Collectively, these results indicate that the overproduction of NO is generally disruptive to locomotor rhythms and suggest that the NO production and clearance should be tightly regulated. LD behavior was not obviously affected in any manipulation (Suppl. Fig. S4).

To find cell types that natively produce and secrete NO and contribute to the control of locomotor rhythms, we next performed an opposite experiment. We expressed RNAi against NOS (VDRC #27725) using a similar set of drivers and analyzed its effects on behavioral rhythms (Table 1). Consistent with the likely absence of NOS within the s-LNvs, NOS RNAi with *Pdf-GAL4* and *R6-GAL4* did not show any behavioral phenotype. NOS RNAi driven with a mushroom body driver *DH52-GAL4* caused a reduced rhythmicity, whereas *macNOS* expression with the same driver had no effect. Most of the other drivers that disrupted rhythms with *macNOS* expression also reduced behavioral rhythmicity with *NOS* RNAi. These include a pan-neuronal driver *R57C10-Gal4*, optic-lobe drivers *79D040-Gal4* and *85B12-Gal4*.The strongest effect was observed with *Repo-GAL4* and *tim-Gal4*, whereas there was no effect with *Clk1982-Gal4*. Since the expression of *Clk1982-Gal4* is relatively restricted to CLK-positive neurons, these results suggest TIM-positive glial cells as an important source of NO in the regulation of circadian locomotion. *Repo-Gal4* driving a second independent RNAi against NOS (TriP #50675) also reduced free-running rhythmicity (3% rhythmic, period 24.0 ± 0 h, N=31). Another VDRC RNAi line (#108433) had no effect on behavior with any driver, which is most likely due to an inefficient knockdown compared to the VDRC #27725 line, judging from the NO staining intensity (Supplementary figure Fig S5). Behavioral patterns in LD was not affected by NOS knockdown in any driver (Suppl. Fig. S4). Altogether, the results of NOS gain -and loss-of-function mini screens indicate that NO produced in many different cell types excluding pacemaker neurons contribute to generate normal free-running locomotor rhythms.

### NO produced in glia plays an active role in the regulation of locomotor output

Constitutive NOS knockdown may induce developmental malformations in the brain that lead to the reduction of rhythmicity, as evidenced by the phenotypes of *NOS Δ* mutants (Figs. 2A, 2B, 3A and 3B). Therefore, to test if NOS is required for active maintenance of rhythmicity after eclosion, we performed the adult-specific knockdown of NOS using pan-neuronal, optic lobe specific and glial GAL4 drivers combined with the temperature-sensitive GAL4 repressor, GAL80^ts^ (49) (Table 1). Glia-specific NOS knockdown caused a notably strong reduction of rhythmicity. In addition, NOS RNAi driven by the pan-neuronal *R57C10-Gal4* extended the free-running period. These results indicate an indispensable role of NO produced in glia for generating circadian locomotor output in adult flies, as well as an existence of a neuronal circuit through which NO signaling regulates the free-running period.

## Discussion

Gaseous signaling molecules play important roles in a myriad of biological processes, including circadian rhythms in mammals. NO and CO were proposed to be “light” and “dark”-induced messengers that convey information from zeitgebers to the core of the molecular clocks (26,50,51). Here we investigated the possible involvement of NO in *Drosophila* circadian rhythms. Our results overall suggests that NO exerts a coordinated, temporarily and spatially diverse effect on the *Drosophila* circadian system.

It is rather surprising that the lack of NOS enzyme is not lethal as NO is part of various developmental processes (18–20,39,52,53). Nonetheless, *NOS Δ* mutants are strongly arrhythmic, incapable of morning anticipation and exhibit minimum light-induced startle response. Whereas the absence of light-induced startle phenotype is likely to be explained by disruptions in light reception through wrong patterning of the receptor cells of the compound eye (39), lack of anticipation point to the problems with the morning pacemakers, the s-LNvs. Indeed, axonal terminals of these neurons in the mutants have an utterly wrong shape, suggesting the wrong or absent synaptic connections with the downstream partners. Together with the axonal morphology phenotype induced by *UNF* knockdown, it is plausible that NO acts through *UNF* and its dimerization partner E75 to define the state of the axonal terminals during development. As in the case of MB neurons (20), this process might involve axon pruning and regrowth.

The functional isoform *dNOS1* showed a circadian variation of its RNA levels throughout the day, which suggest that levels of NO could cycle. However, *dNOS* is likely to be regulated by its truncated isoforms in a stage -and cell-type-specific manner, which lays an additional complexity to the regulation of NO production and probably leads to the heterogeneous and context-specific variations of NO. Whether its cycling is important or not for the rhythmicity, it is clear that NO has only a modulatory role in the s-LNvs’ molecular clocks, since UNF or E75 knockdown in the s-LNvs have much stronger effect on molecular clockwork, i.e. dampening of the PER cycling amplitude and period extension (14,12).

In our *NOS-RNAi* mini screen, two non-clock neurons-specific drivers, *GMR79D04* and *GMR85B12* reduced locomotor rhythmicity in DD. This phenotype was rather specific to developmental stage, reinforcing the idea that NO is necessary for a proper establishment of neuronal circuits. A low rhythmicity phenotype produced by knockdown under pan-neuronal driver *GMR57C10-GAL4* backs up this idea. Intriguingly, however, in addition to the low rhythmicity, *GMR57C10 > NOS RNAi* in adulthood resulted in an extended period. This raises a question on what might be the neuronal subsets that produce NO and contribute to the regulation of period length of locomotor activity. The effect must be of a different nature from the regulation of *per* transcription by UNF/E75. An interesting hint comes from a recent study by the group of G. Rubin (54), which shows that NO acts as a co-transmitter in a subset of dopaminergic neurons, specifically in some of the PAMs, PPL1s and PPL2abs. It is thus possible that dopamine signaling regulated by NO is involved in the control of locomotor activity period.

Targeting glial cells leads to the strongest and most persistent phenotype in locomotor activity both for gain -and loss-of-function of NOS. The importance of glia in circadian rhythms have been recognized, especially those expressing the molecular clocks and exert reciprocal communication with the pacemaker neural circuit (55,56). Our study is the first to identify NO as a signaling molecule produced in glia that mediates part of the role of glia in circadian behavioral in flies. It has been shown that in mammals NO mediates light-induced phase-shifts through the cGMP pathway (31). It is an interesting parallel to note that forced production of NO in the s-LNvs caused phase shift rather than amplitude dampening, although we speculate that part of this effect comes from NO hampering E75/UNF dimerization that normally enhances CLK/CYC-mediated *per* transcription. Mammalian clocks contain E75 homologs REV-ERB *α/β* that work with another nuclear receptor, ROR. In contrast to flies, REV-ERB *α/β* have a repressive function within the clocks, inhibiting transcription of *BMAL1*, a mammalian analog of *CYC*. Interestingly, *in vitro* studies of mammalian cell culture showed that excessive presence of NO increases the production of *BMAL1*, consistently with the hypothesis that NO decreases REV-ERB *α/β* activity (57). These findings altogether point out that NO is an evolutionarily conserved regulator of circadian rhythms.

In line with recent studies (7,58,59), our research expands the view on the factors that participate in neuronal and molecular mechanisms of circadian rhythmicity. The finding that gaseous messenger NO contributes to the various aspects of circadian rhythmicity, from development to the maintenance, emphasizes that non-cell-autonomous, systemic regulation is integral to the circadian circuit operation. Our results set a foundation for future studies addressing whether or not specific glial or neuronal classes are required in this regulation and how the NO signaling modulates the state of the pacemaker circuit.

## Materials and Methods

### Fly rearing, crosses, and strains

*Drosophila* were reared at 25 °C on a corn-meal medium under 12 hr:12 hr light-dark (LD) cycles. Two CRISPR deletion mutants *NOS Δ ter, NOS*Δ*all* were kindly provided by O. Schuldiner (20). The *UAS-macNOS* line was originally generated by H. Krause (18) and provided also by O. Schuldiner. The drivers *GMR57C10, GMR79D04, GMR85B12, GMR33H10* (60) as well as *UAS-NOS-RNAi*^*56675*^ were obtained from Bloomington Stock Center (Indiana, US). The UAS lines *NOS-RNAi*^*27725*^ and *NOS-RNAi*^*108433*^ were obtained from the Vienna *Drosophila* Resource Centre (VDRC). The *Clk1982-Gal4* line was provided by N.R. Glossop (61). The lines *Pdf-Gal4* (62), *Repo-Gal4* (63), *OK107-Gal4* (64), *D52H-Gal4* (65), *GMR-Gal4* (66), *Elav-Gal4* (67), *R6-Gal4* (68), *UAS-miR unf* (12,14) were described previously.

### Behavioral Assays

The locomotor behavior assay was performed as described previously (12) and data were analyzed using FaasX software (69). Briefly, male flies were first entrained in 12 h/12 h LD cycles for 4 days and then released in DD for 7–10 days. The flies with power over 20 and width over 2.5h according to the χ^2^ periodogram analysis were defined as rhythmic. The significance threshold was set to 5%. The χ^2^ test was used to compare the rhythmicity of the flies, and the Student’s *t* test (2-tailed) was used to compare the free-running period.

### Immunocytochemistry, microscopy and quantification

The brains were imaged using a Leica SP5 confocal microscope. At least 10 brain hemispheres were subjected to analysis using Image J software (National Institutes of Health). The anti-PER signal was quantified as previously described (12).

### Nitric oxide visualization and measurements

NO visualization was performed as described in (20) with minor modifications. Brains were dissected in PBS and incubated with 10 μM Diaminorhodamine-4M AM (DAR-4M, Sigma-Aldrich) in PBS for 1 h at RT, followed by the fixation for 15 min in PBS containing 4% paraformaldehyde. Immediately after the fixation brains were mounted and imaged. For NO measurements at different times of the day, the procedure was exactly the same with the omission of the fixation step. Long-term NO measurement in *ex-vivo* brain culture was performed as described in (70). Briefly, brains were dissected on an ice-cold plate in modified Schneider’s medium (SMactive) (71) with an addition of 5 mM Bis-Tris (Sigma) and then mounted on a glass-bottom dish (35 mm MatTek petri dish, 20 mm microwell with 0.16/0.19 mm coverglass). The glass-bottom well was filled with the SMactive medium with 10 μM DAR-4M. Time-lapse imaging was performed at 25 °C and 80%, with images acquired every hour.

### RNA analysis

Total RNA was isolated from adult fly heads using Trizol (Invitrogen) following the manufacturer’s protocol. The RNA was reverse-transcribed using oligo(dT) primers, and the resulting cDNAs were quantified using real-time qPCR as previously described (47). The mRNA levels of *dNOS1* were normalized to those of *elongation factor 1 (Ef1).*

## Supporting information

Supplementary Fgure S5

Supplementary Fgure S3

Supplementary Fgure S2

Supplementary Fgure S1

Supplementary Fgure S4

## Acknowledgments

We thank O. Schuldiner and N. Glossop, the Bloomington Drosophila Stock Center and Vienna Drosophila Resource Center for fly stocks. We are grateful for our lab members for valuable discussions on this work.

## Supporting Information Captions

**Supplementary Figure 1. NOS splice isoforms.** dNOS1 and presumably dNOS8 encode functional complete enzyme. The rest isoforms lead to a truncated enzyme. Information is taken from (34). dNOS1-specific primers used are (F) GGC GAG CTT TTC TCC CAG GA, and (R) GAC GAG CCA ATG CTG GAG TC, indicated in red.

**Supplementary Figure 2. Axonal morphogenesis of the s-LNvs relies on UNF.** s-LNv axonal terminals in control (*Pdf-GAL4*/+) and flies with LNv-targeted UNF knockdown (*Pdf-GAL4* driving *UAS-miR unf*, shown as *PDF> UNF-RNAi*) were visualized by co-expressing *mCD8::Venus*. Flies were dissected at ZT2 and stained with anti-GFP antibodies. Representative confocal images are shown.

**Supplementary Figure 3. Upregulation of NO upon macNOS overexpression.** DAR4-M staining of brain expressing *macNOS* in *Pdf-Gal4*. Left, Representative confocal images. Right, comparison of DAR4-M fluorescent levels of a single representative WT and *Pdf* > *macNOS* brains within the region of s-LNvs.

**Supplementary Figure 4. Cell-restricted manipulation of NOS does not affect LD locomotor behavior.** Locomotor activity histograms for group average activity of macNOS or NOS-RNAi^27725^ expressed under indicated drivers. 4 days of LD are shown.

**Supplementary Figure 5 NOS-RNAi efficiency comparison.** DAR4-M staining measures of two *Elav-*driven NOS-RNAi^27725^ and NOS-RNAi^108433^. Fluorescence levels were measured broadly in the region of the central brain, approximately in the area of the central complex. RNAi line 27725 induces a significant reduction of DAR4-M signal. *\*\*p*<0.01 (Student’s test).

## Tables

**Supplementary Table S1.** Optic lobe-specific drivers. Distribution and intensity of the tested generic optic lobe-specific drivers, taken from Janelia Fly Light project. Original characterization was based on the GFP expression. All three optic lobe-specific drivers are enriched in the optic lobes. Occasional expression outside OL is mainly relatively weak in compare with expression in OL.

|  |  |  | R33H10 | R79D04 | R85B12 | R57C10 |
| --- | --- | --- | --- | --- | --- | --- |
| antennal lobe |  |  | 1_2 | 1_1 | 1_5 |  |
| antennal mechanosensory and motor center |  |  | 1_3 | 0_5 | 5_0 |  |
| anterior ventrolateral protocerebrum |  |  | 3_0 | 2_1 | 1_1 |  |
| antler |  |  | 1_1 | 1_0 | 0_5 |  |
| bulb |  |  | 1_4 | 1_0 | 1_0 |  |
| cantle |  |  | 1_2 | 0_5 | 1_0 |  |
| crepine |  |  | 1_1 | 0_5 | 1_1 |  |
| ellipsoid body |  |  | 0_5 | 0_5 | 0_5 |  |
| epaulette |  |  | 1_4 | 2_2 | 1_1 |  |
| fan-shaped body |  |  | 1_1 | 2_0 | 1_0 |  |
| flange |  |  | 2_1 | 2_1 | 2_2 |  |
| gall |  |  | 1_2 | 0_5 | 0_5 |  |
| gorget |  |  | 1_2 | 3_1 | 1_1 |  |
| inferior bridge |  |  | 1_1 | 1_0 | 1_0 |  |
| inferior clamp |  |  | 1_2 | 1_1 | 1_1 |  |
| inferior posterior slope |  |  | 1_2 | 1_0 | 1_1 |  |
| lateral accessory lobe |  |  | 1_2 | 3_0 | 1_1 |  |
| lateral horn |  |  | 1_1 | 1_2 | 1_0 |  |
| mushroom body |  |  | 1_1 | 1_0 | 1_2 |  |
| noduli |  |  | 0_5 | 0_5 | 0_5 |  |
| optic lobe |  |  | 5_2 | 5_2 | 5_2 |  |
| optic tubercle |  |  | 2_0 | 0_5 | 2_0 |  |
| posterior lateral protocerebrum |  |  | 1_2 | 2_1 | 1_0 |  |
| posterior ventrolateral protocerebrum |  |  | 2_1 | 1_3 | 1_2 |  |
| protocerebral bridge |  |  | 0_5 | 1_0 | 1_0 |  |
| prow |  |  | 1_4 | 2_3 | 4_1 |  |
| saddle |  |  | 2_1 | 1_0 | 4_0 |  |
| subesophageal ganglion |  |  | 3_2 | 3_0 | 2_1 |  |
| superior clamp |  |  | 2_1 | 1_1 | 1_0 |  |
| superior intermediate protocerebrum |  |  | 2_1 | 1_0 | 1_1 |  |
| superior lateral protocerebrum |  |  | 1_2 | 1_0 | 2_0 |  |
| superior medial protocerebrum |  |  | 2_1 | 1_1 | 1_2 |  |
| superior posterior slope |  |  | 1_1 | 1_1 | 0_5 |  |
| vesta |  |  | 1_4 | 0_5 | 1_0 |  |
| Gene region |  |  | Src64b | CG9650 | zf30c | Nsyb |
| Intensity |  |  |  | Distribution |  |  |
| Value | Description | Criteria for selection |  | Value | Description | Criteria for selection |
| 0 | blank | No pattern |  | 0 | passing tract | Tracts passing neuropil |
| 1 | faint | Visible with enhanced contrast only |  | 1 | sparse | Like one glom in al |
| 2 | very weak | Visible when known with nc82 staining |  | 2 | local, regional | Bigger region or multiple glom in al |
| 3 | weak | Easily visible with nc82 staining |  | 3 | widespread | Less than half of the structure |
| 4 | strong | Bright |  | 4 | prevalent | More than half of the structure |
| 5 | very strong | Really bright |  | 5 | ubiquitous | Everywhere |

