## Supplementary figures and images for "Nitric Oxide Mediates Neuro-Glial Interaction that Shapes *Drosophila* Circadian Behavior"

### Supplementary Fgure S1

#### Supplementary Figure 1

##### NO Synthase Splice Variants

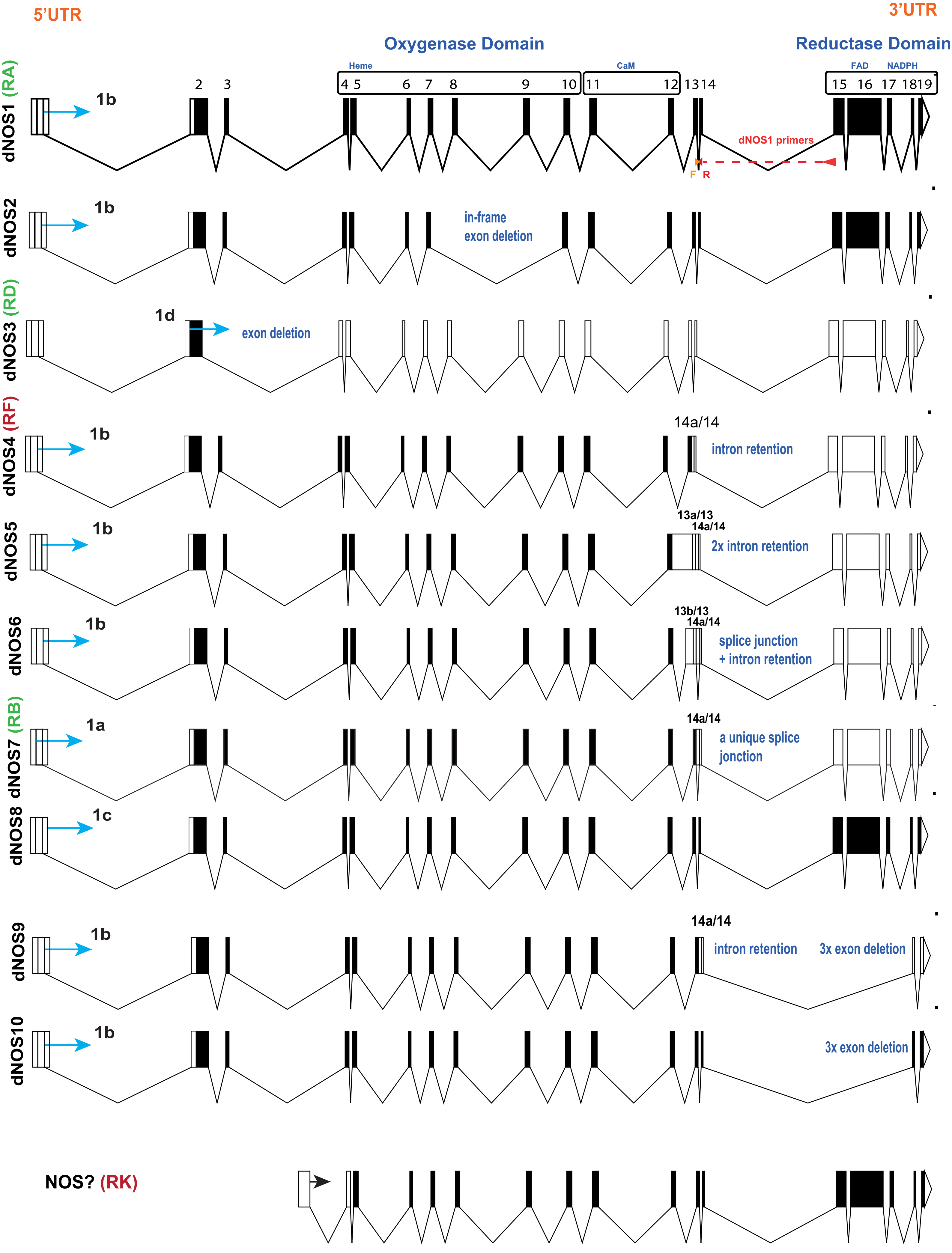

### Supplementary Fgure S2

#### Supplementary Figure 2

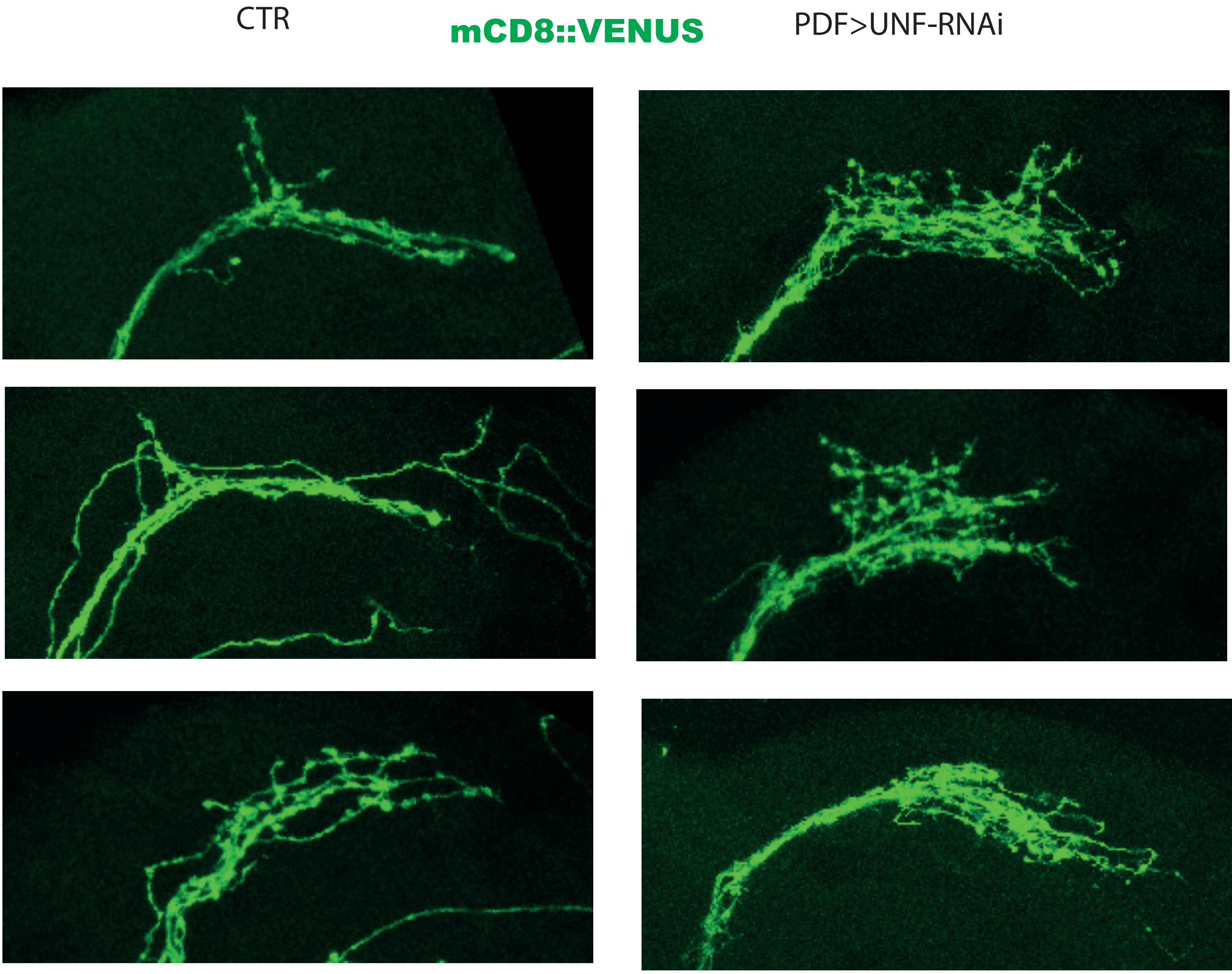

### Supplementary Fgure S3

#### Supplementary Figure 3

PDF>macNOS

**DAR4-M, LNVs**

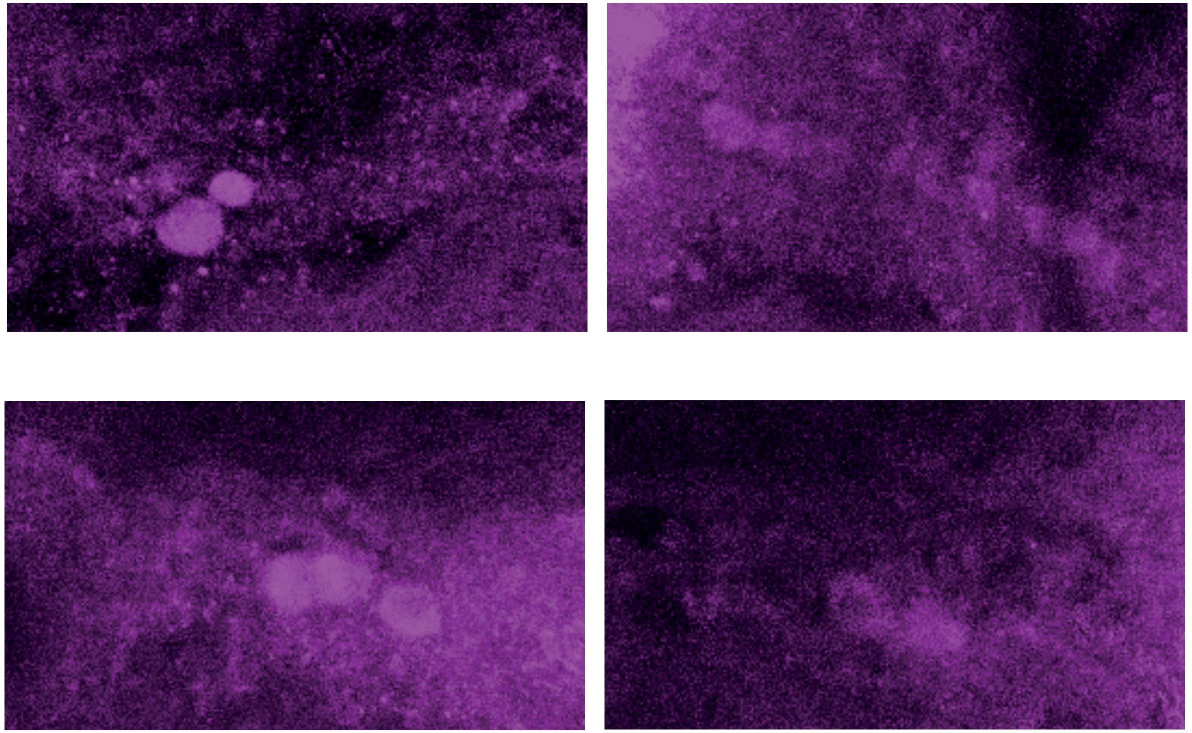

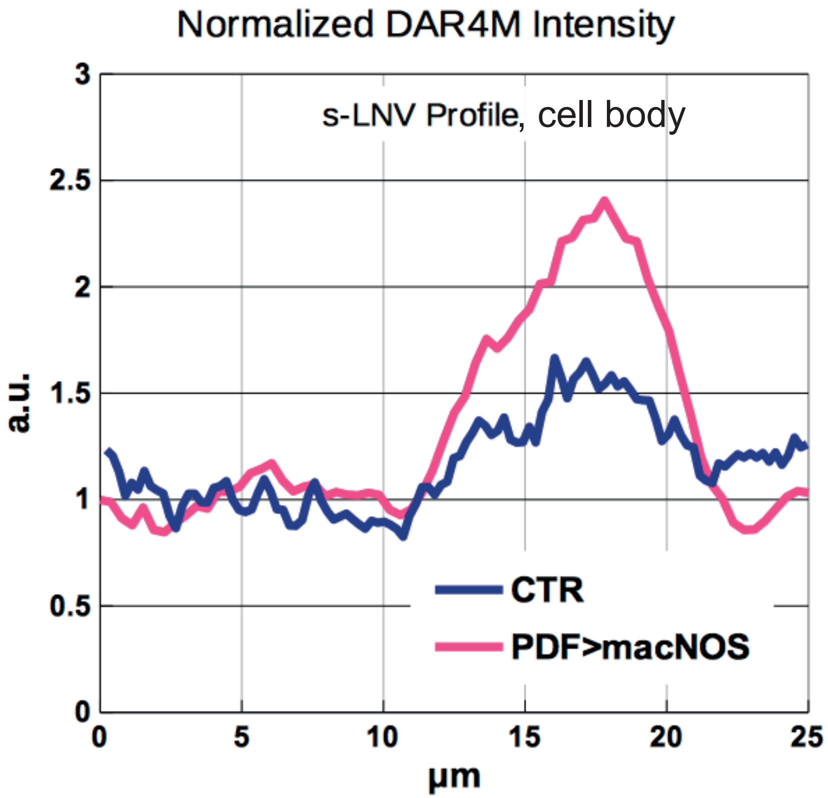

### Supplementary Fgure S4

Supplementary Figure 4

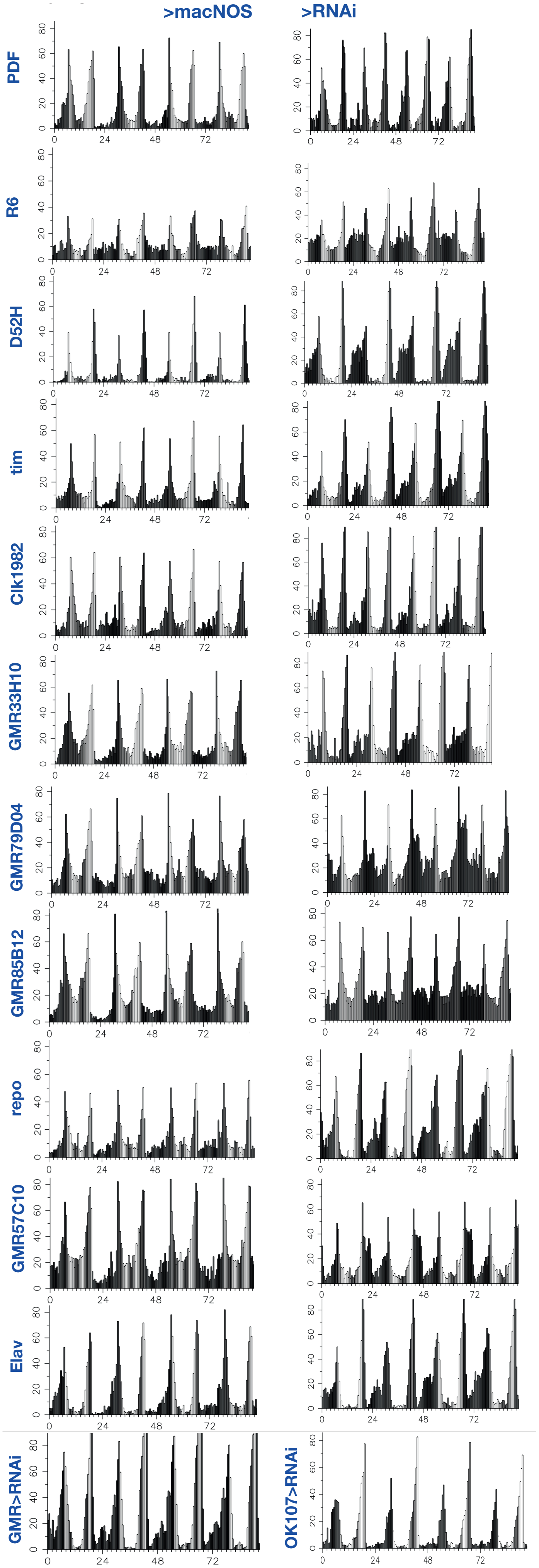

### Supplementary Fgure S5

#### Supplementary Figure 5

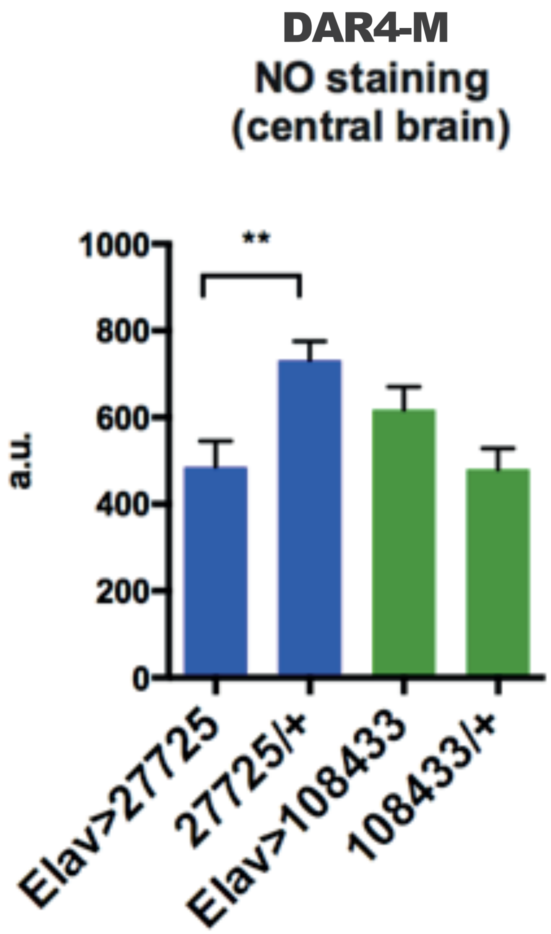
